# Hardware evaluation of spike detection algorithms towards wireless brain machine interfaces

**DOI:** 10.1101/2022.06.23.497414

**Authors:** Alexandru Oprea, Zheng Zhang, Timothy G. Constandinou

## Abstract

The current trend for implantable Brain Machine Interfaces (BMIs) is to increase the channel count, towards next generation devices that improve on information transfer rate. This however increases the raw data bandwidth for wired or wireless systems that ultimately impacts the power budget (and thermal dissipation). On-implant feature extraction and/or compression are therefore becoming essential to reduce the data rate, however the processing power is of concern. One common feature extraction technique for intracortical BMIs is spike detection. In this work, we have empirically compared the performance, resource utilization, and power consumption of three hardware efficient spike emphasizers, Non-linear Energy Operator (NEO), Amplitude Slope Operator (ASO) and Energy of Derivative (ED), and two common statistical thresholding mechanisms (using mean or median). We also propose a novel median approximation to address the issue of the median operator not being hardware-efficient to implement. These have all been implemented and evaluated on reconfigurable hardware (FPGA) to estimate their hardware efficiency in an ultimate ASIC design. Our results suggest that ED with average thresholding provides the most hardware efficient (low power/resource) choice, while using median has the advantage of improved detection accuracy and higher robustness on threshold multiplier settings. This work is significant because it is the first to implement and compare the hardware and algorithm trade-offs that have to be made before translating the algorithms into hardware instances to design wireless implantable BMIs.

## I. Introduction

BMIs have enabled disabled patients to control neuroprosthetics [1] or communicate through thought [2] [3]. In such circumstances, intracortical implants record the electrical activity of the neurons and subsequently analyze the spikes (observed action potentials) through a behavioral decoding algorithm in order to control external assistive technology such as a prosthetic limb or a text-to-speech prosthesis.

With the recent trend of designing wireless distributed BMIs with increased channel counts [4], the growing amount of data requires unfeasible bandwidth and wireless transmission power for the implants. Real-time on-implant feature extraction and compression are therefore essential to distill the informative features, reduce bandwidth and consequently reduce the transmitting power. The firing rate is one of the most commonly used features [5]–[8] representing essential information correlated to subject behaviors with significantly reduced bandwidth. Using firing rate reduces the bit rate linearly with the increase of bin count period [4]. However, on-implant feature extraction and compression bring extra hardware complexity and face great challenges in power consumption. It has been demonstrated in [9] that in practice, the implant power consumption should be less than 0.8 mW/mm^2^.

To obtain the firing rate of neurons, the neuron action potential spikes have to be detected and counted. The detection of spikes represents the process of discerning between the background noise of the signal and the action potential. Multiple spike detection techniques have been developed based on template matching [10] [11], wavelet transform [12], and statistics [13]–[15]. Due to its simplicity, the latter class of methods has generated substantial interest in fully-implantable BMIs. The expectation is generally the most used statistics due to its low implementation cost in the form of a moving average. The mean is usually paired with the NEO or its variants (i.e., ASO, ED, Smoothed-NEO, multi-NEO) [16]–[19] which emphasizes the spikes before averaging. The median is regarded in the literature as a measure of the standard deviation of the noise through the sensible assumption of normal distribution [20]. However, the successive comparison and large data buffer involved make it less feasible in hardware use. Similarly, due to high hardware cost, other thresholding mechanisms using statistics such as standard deviation, RMS value, and cross-correlation [15] have also not been widely used in hardware implementations.

There are significant studies on implementing and comparing various spike detection algorithms, but they rarely experimentally compare their hardware trade-offs. In this work, we have implemented three different spike emphasizers (NEO, ASO, ED), two thresholding algorithms (mean and median), and a compression module (spike binner). We have also proposed a novel implementation of median thresholding to fit the median operation in an allowable hardware budget. A comprehensive comparison is drawn in terms of spike detection performance of algorithm combinations and Field Programmable Gate Arrays (FPGA) power consumption and resource usage.

## II. System Architecture

Fig. 1 depicts the general signal path of a fully-implantable BMI with on-implant signal processing capability. This work will focus on the last three phases (i.e., Emphasizers, Thresholding, and Compression). Our implementation assumes an analog fronted of 7 kHz and 10-bit resolution per channel, as they have been proved to be the minimum requirements to contain the essential information of the spikes [21]. As this literature also suggested, a two-pole Butterworth filter is used to remove the LPF leaked to the passband.

**Fig. 1.**
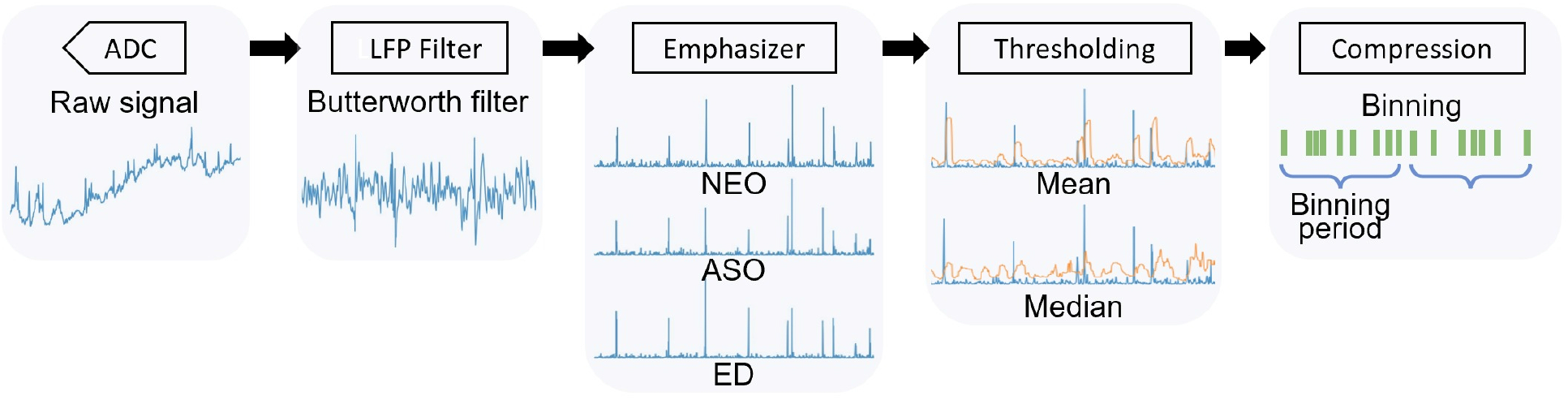
The architecture of the firing rate-based BMI. The waveform under each module represents the output wave of the respective stage

### A. Emphasizer

Before finding the threshold, a pre-emphasis step is applied to the LPF removed signal. Traditionally, NEO is used due to its characteristics of accentuating high energy, high-frequency signals (i.e., spikes) to suppress the noise.

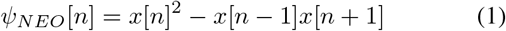

Eq. 1 depicts the formula of the digital NEO operator *ψ*_*NEO*_. There are multiple variants of NEO; however, some involve digital filter smoothing [18] or more samples to be buffered [19], adding extra complexity. Therefore, the only variants of NEO considered in this work are ASO [16] and ED [17], due to their low hardware cost, requiring one less multiplication and one less sample than the NEO. ASO and ED are shown in formula 2 and 3 respectively.

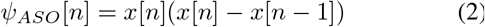

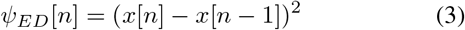

After the computation of the emphasized signal, the absolute values are then taken to preserve the amplitude and gradients on both sides.

### B. Thresholding Algorithm

Subsequent to pre-emphasis, the signal is processed with the thresholding module. Traditionally, after the NEO pre-emphasis, the threshold would be set using a moving average as in the formula below.

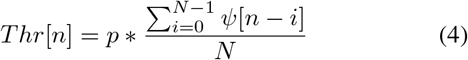

Where N is the current sample, *ψ* is the output of the emphasizer, *p* is a constant, and N is the number of samples included in the average. This solution is employed due to its very low computational cost as on every cycle, an addition, a subtraction, and a shift is performed (Given that *p* and N are carefully chosen). This method was proved to yield good performance [13] considering its simplicity. N here is set to be 16, yielding good performance, low resource occupation, and ease of bit shift.

The median operation has a high implementation complexity of over O(*n*^2^), which is unsuitable for real-time processing. In order to mitigate these shortcomings, we propose to successively find the median of medians within a set of samples to estimate the median values. The approach divides the sequence into sets of 5 and then finds the median of each set. The process then repeats to the medians found until only one number remains. This process does not guarantee to find the median but an estimation. However, our results suggest that the accuracy drop is under 3% in low to moderate noise scenarios, reaching approximately 5% in high noise ones (Fig. 3).

This approach represents a hardware-efficient implementation as the implementation complexity has decreased from over O(n^2^) to O(nlog(n)). Furthermore, the instances of the modules can be set up through simple interconnections, allowing for scalability. The set size of 5 was chosen as the minimum interval that can contain a spike by itself. A higher interval would have generated a behavior closer to a median. However, it would have come at the cost of multiple successive comparisons. The implementation of the median of 5 number module achieves the optimal resource consumption by using only 6 comparators. Notice that the median mentioned later is this median approximation using 25 samples unless specified.

### C. Compression

The compression module is constructed as a series of counters (one for each channel) that count the number of threshold crossing in every 800 samples (about 114 ms). To allow more robustness in the binning process and avoid multiple detections of one spike, the binner will not increment the value after one detection, until one refractory period has passed, usually 1 ms. At the end of the set binning period, counts are output, and the counters reset.

## III. Results

### A. Evaluation metrics and hardware platform setting

To test the spike detection performance of the algorithms in combination with the different emphasizers, the algorithms have been simulated offline on a widely used dataset from [20]. It consists of real spike shapes positioned in time with Poisson distribution. The superimposed realistic noise consists of randomly selected spikes with a relative standard deviation of 0.05, 0.1, 0.15 and 0.2 respectively. Accuracy, a generally used metric to compare spike detection algorithms, is presented below.

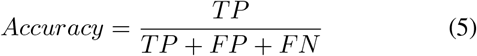

Where True Positive (TP) represents truly detected spikes, False Positive (FP) represents wrongly detected spikes, and False Negative (FN) represents the undetected spikes.

Different methods have been implemented on the XC7A35T AMD FPGA built on the 28nm technology. The resource utilization was reported by the synthesizer and the dynamic power was measure by subtracting between the FPGA core power with and without the methods implemented. In order to avoid simplification in the synthesis stage, the out-of-context mode is used. These steps assure that the reported power is directly linked to the complexity of the algorithm and not to synthesis factors. The FPGA hardware results would not translate directly to the ultimate ASIC design, but it is still a reliable estimation for the ease of algorithm complexity comparison and implementation selection.

### B. Algorithm performance

From Fig.2 (a), out of the three emphasizers, ED (Solid lines) yields the highest accuracy across all noise levels and with both methods. ED also exhibits the highest adaptability through robustness against multiplier choice (flatter curves). ASO (Dotted lines) shows similar characteristics but reaches lower accuracy at higher noise levels. The performance of NEO (Dash lines) deteriorates rapidly as the level of noise increases. This finding suggests that the gradient is more discriminative than amplitude in spike emphasizing, especially when noise increases, ED outstands.

As for median approximation (MA, Red lines) and average (A, Blue lines), their performance is similar w.r.t the highest accuracy they can achieve in different noise levels as the figure shows. However, it is a common drawback of the statistical-based thresholding that multiplier choosing can affect the detection performance [22] and therefore introduces the effort of manually tuning the threshold in practice. The median, as the figure shows, is more robust to the choice of the multiplier as it generates a flatter response close to its maximum point. This behavior is most noticeable at 0.05 noise, where the algorithm can always yield high accuracy as long as the multiplier is large enough. On the other hand, when using the average, the accuracy decays very fast as the multiplier gets further from the peak values. This susceptibility is mitigated by using ED, while it results in the most narrow interval by using NEO. Furthermore, The interval for choosing a multiplier narrows as the noise increases for the median method, however, its effect is less noticeable across all emphasizer choices.

Based on the findings above,using ED alongside median approximation can provide the highest spike detection performance while it is the least affected by the choice of the multiplier.

### C. The effect of the number of samples

We also assessed how the number of samples used to calculate the median and average affects the detection performance in high noise level cases. Detecting spikes at high noise levels can be challenging as more numbers need to be buffered for mean/median calculation, but the number buffering can be resource-consuming.

The average starts outperforming the median method as the noise increases with 16 samples for the mean and 25 samples for the median. However, suppose we allow more samples to use. In that case, when the number of samples is increased to 90 samples for the average and 50 for the median, the detection accuracy can be improved by 5%, and the median outperforms the average at all noise levels; Fig. 2(b) shows this behavior. Therefore, if the available memory is increased, the median yields both higher accuracy and better robustness.

**Fig. 2.**
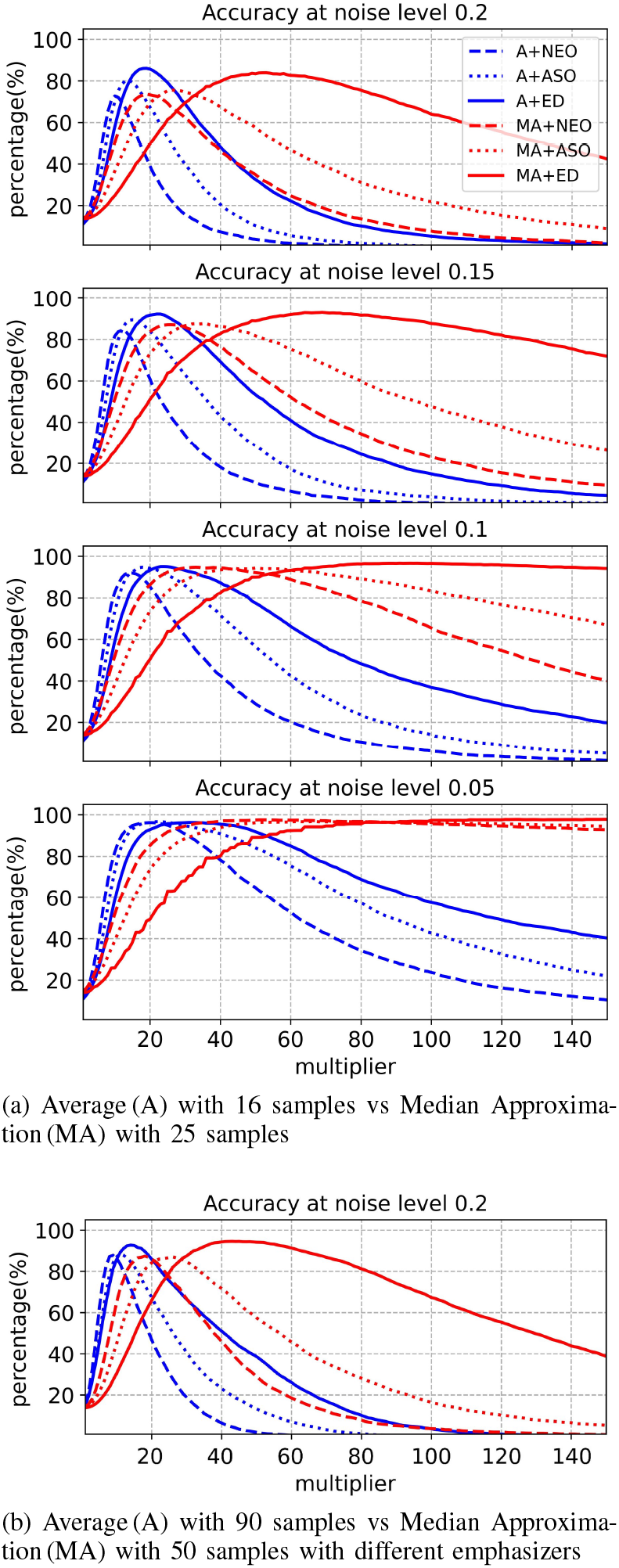
The performance difference of Average (A) vs. Median approximation (MA)

**Fig. 3.**
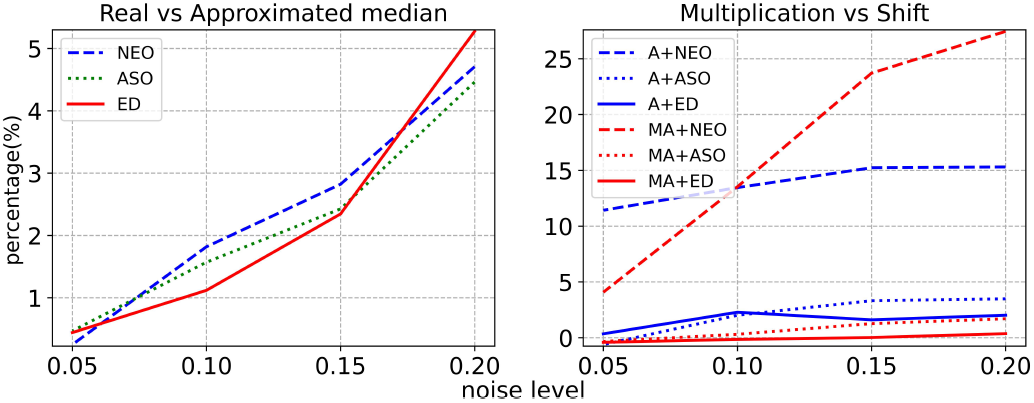
Accuracy degradation of approximations median (left) and multiplication (right)

### D. Resource consumption

One of the most costly hardware resources in the presented emphasizers is the multiplier. They was implemented in three methods: Look-Up Tables (LUTs), Digital Signal Processing units (DSP), and an approximation by left shifting according to the index of the most significant bit. The latter has been previously employed in spike detection yielding promising results [16]. Table. I shows bit shift approximation uses half the resources of the LUT-based multiplier representing a reliable comparison to their ASIC implementation. However, the shift approximation is unsuitable when using the NEO emphasizer as the accuracy degradation overpasses 10% shown in Fig.3. The other two emphasizers exhibit a 4% degradation at worst. The median with ED, in particular, exhibits almost no accuracy degradation. Using DSP yields the lowest power consumption as it is customized for such operation, but DSP would occupy the extra area in ASIC design. Only the NEO emphasizer was used to compare the multiplier methods because it would indicate a larger power difference. The rest of the emphasizers are implemented using DSPs.

**TABLE I.** The power and resource utilization of different modules

| Module name | LUTs | Registers | Block RAM | DSPs | Power($\mu W$) | Power per channel ($\mu W$) |
| --- | --- | --- | --- | --- | --- | --- |
| Butterworth filter | 196 | 0 | 1 | 0 | 71.8 | 0.75 |
| NEO multiplier | 235 | 0 | 1 | 0 | 58.6 | 0.61 |
| NEO bitwise shift | 120 | 0 | 1 | 0 | 53.0 | 0.55 |
| NEO DSP | 30 | 0 | 1 | 2 | 46.6 | 0.49 |
| ASO DSP | 40 | 0 | 0.5 | 1 | 40.0 | 0.42 |
| ED DSP | 13 | 0 | 0.5 | 1 | 36.4 | 0.38 |
| Average | 41 | 0 | 5 | 0 | 461.8 | 4.81 |
| Median unrolled | 1495 | 0 | 7 | 0 | 634.6 | 6.61 |
| Median rolled | 465 | 184 | 7 | 0 | 1107.4 | 11.54 |
| Counter | 85 | 32 | 0.5 | 0.5 | 25.2 | 0.26 |

The ED uses the least resources and power compared to the other methods as expected. ASO and ED use half the DSP units and the memory used by NEO. ASO uses, unexpectedly, more LUTs than NEO, because the subtraction is fused with multiplication in a single multiply and accumulate (MAC) operation in the DSP core.

The average is implemented through a moving average; thus, it uses few resources outside of the memory. The ram blocks hold both previous samples and the previous value of the average. For our median algorithm, we tested two implementations: one consists of six median modules that divide the 25 samples into groups of 5, and then the median of medians is taken (unrolled version), and the other one has one module fed with 5 groups of five samples and, subsequently, with the medians (rolled version). The first solution aims to reduce the dynamic power consumption at the cost of resource usage, while the other targets the opposite effect. The rolled version uses approximately a third of the logic units used by the unrolled version. However, this improvement comes at the cost of registers that feed the median module and a substantially increased power consumption. The steep increase in state transitions causes the power to almost double in the case of the rolled version of the median.

The counter module uses memory to store the number of arrival spikes in each channel and the bin period counts. The output remains unchanged until the bin period is reached. Due to the low output frequency, it has a lower power consumption than other modules using similar resources (e.g., emphasizers).

## IV. Conclusion

This paper discusses the trade-off between hardware complexity and detection performance of the median approximation and moving average preceded by three hardware-efficient emphasizers (NEO, ASO, ED) for spike detection. Multiple considerations were taken into account, such as the number of samples, different multiplication methods, and median module reusing.

ED demonstrated superior performance with both thresholding algorithms and used fewer resources. It not only reaches a larger peak accuracy but also enables adaptability to multiple noise levels. On the other hand, the proposed median approximation makes the median operation achievable in hardware with an acceptable budget and easily scalable. Using the median approximation consumes more power and resources than the average thresholding, but it results in better spike detection performance and improved robustness to signal variations. At low noise levels, the median approach does not require tuning in multiplier choice, regardless of which emphasizer is used.

